# Disrupting astrocyte signalling in the nucleus accumbens impairs incentive-driven instrumental actions

**DOI:** 10.64898/2026.01.12.699167

**Authors:** Joanne M. Gladding, Octavia Soegyono, Arvie Rodriguez Abiero, Karly M. Turner, Michael D. Kendig, Laura A. Bradfield

## Abstract

Astrocytes in the nucleus accumbens (NAC) core have been observed to undergo phenotypic changes associated with drug-seeking behaviour in both humans and animals. However, the role of NAC core astrocytes in non-drug-related instrumental behaviour remains poorly understood. To address this, we chemogenetically activated hM4Di receptors selectively expressed on NAC core astrocytes in rats during food-motivated decision-making tasks. In Experiment 1, rats were first trained to associate two auditory stimuli with two distinct food outcomes (pellets and sucrose), then to press left and right levers for those same outcomes. All training was conducted drug-free, and rats then received intraperitoneal (i.p) injections of either vehicle or deschloroclozapine (DCZ) prior to test. Disrupting astrocytic signalling via DCZ injections left instrumental choice intact when it was guided by cues signalling the sensory-specific properties of each outcome, as tested in specific Pavlovian instrumental transfer and outcome-selective reinstatement, but suppressed responding in an outcome devaluation test. In Experiment 2, a single stimulus and single lever were separately paired with distinct food outcomes, then presented together on test. Control animals demonstrated a general PIT effect, elevating responding during stimulus presentations, and this was prevented by Gi activation on NAC core astrocytes. Immunohistochemistry revealed increased neuronal activity following hM4Di activation in astrocytes. Together, these findings suggest that intact signalling in NAC core astrocytes is necessary for instrumental actions that depend on general arousal or affective processes, but not for actions guided by sensory-specific outcome expectations.

**Significance Statement:** Astrocyte dysfunction in the nucleus accumbens (NAC) has been implicated in several forms of compulsion in humans, yet preclinical work has focussed almost exclusively on drug-taking and seeking. The role of NAC core astrocytes on responding for non-drug outcomes is therefore unclear. Here, using food outcomes, we show that disrupting astrocytic Gi signalling in NAC core selectively impairs actions driven by general motivational states but spares those guided by specific outcome expectations. These findings suggest that NAC core astrocytes play a critical role in invigorating behaviour, extending their involvement beyond drug-seeking in animals and highlighting their potential relevance to compulsive behaviour more generally.

## Introduction

Decision-making is influenced by environmental cues that guide actions towards motivationally relevant outcomes (Doya, 2008). This capacity for cues to motivate behaviour has been shown to influence the development and persistence of addiction and other compulsive behaviours (Garbusow et al., 2016; Krypotos & Engelhard, 2020; Peng et al., 2022). In particular, in rats compulsive drug-seeking behaviours have been linked to enhanced sensitivity to drug-related cues and reduced astroglial glutamatergic transport in the nucleus accumbens (NAC) core (Bobadilla et al., 2017; Kalivas, 2009). However, less is known about the role of NAC core astrocytes in naturalistic (e.g. food-driven) decision-making tasks.

Across both rodents and humans, the influence of predictive cues on behaviour has been investigated using Pavlovian-instrumental transfer protocols (Cartoni et al., 2016; Colwill & Rescorla, 1988; Holmes et al., 2010). This procedure involves separate Pavlovian (cue-outcome) and instrumental (response-outcome) learning stages, and the transfer between them is tested when the cues and responses are presented together for the first time. The typical finding, at least for appetitive paradigms, is that the presence of the cue(s) enhances the performance of the action. There are two forms of transfer: general, in which cues induce general motivational arousal, and outcome-specific, in which cues bias responding towards responses associated with specific outcomes.

At the neural level, prior research has shown that excitotoxic lesions of the NAC core given to rats prior to training, and pharmacological inactivation prior to test, attenuates the general excitatory effects of reward-related cues in general Pavlovian-instrumental transfer whilst leaving outcome-specific transfer intact (Corbit & Balleine, 2011; Corbit et al., 2016). Outcome-specific transfer was likewise preserved following neuroinflammation in the NAC core, along with outcome-selective reinstatement, whereas the incentive-value driven process of outcome devaluation was attenuated (Abiero et al., 2025). In that same study, NAC core neuroinflammation also enhanced sensitivity to food-related cues, causing rats to increase the number of head entries they made into the magazine in which food was delivered. In the dorsomedial striatum, the effects of neuroinflammation were linked specifically to the disruption of astrocytic signalling, but no such link has been investigated for the NAC core. The present study therefore examined whether astrocytes in the NAc core regulate responding in food-motivated decision-making tasks. In astrocytes, chemogenetic activation of Gi-protein–coupled receptors does not inhibit activity, as it does in neurons, but instead disrupts normal physiological signalling (Durkee et al., 2019). Accordingly, in the present study we used this approach to test whether disrupting astrocytic signalling in the NAc core alters goal-directed and/or cue-driven action performance.

## Materials and methods

### Experiment 1. The effect of disrupted NAC core astrocyte signalling on specific PIT

#### Animals

A total of 38 Long Evans rats (18 female and 20 male) were purchased from Ozgene (Perth, Australia). Three groups were generated: hM4Di+VEH (*n* = 14, 7 female, 7 male), mCherry+DCZ (*n* = 11, 6 female, 5 male), and hM4Di+DCZ (*n* = 13, 5 female, 8 male). Rats were at least 10 weeks old at the beginning of the experiment and housed in transparent plastic boxes (up to 3 rats per box) in a temperature- and humidity-controlled colony room maintained on a 12h light-dark cycle (lights on between 7:00am-7:00pm). All behavioural procedures were conducted during the light cycle. Water and standard lab chow were available *ad libitum* prior to the start of each experiment. During behavioural training and testing, rats were maintained at ∼90% of their *ad libitum* body weight. All procedures were approved by the Animal Care and Ethics Committee at the University of Technology Sydney (ETH21-6657).

#### Apparatus

Training and testing took place in 14 identical Med Associates (Fairfax, VT, USA) operant chambers enclosed in light- and sound-attenuating cabinet. Each chamber contained a recessed food magazine connected to a pellet dispenser (45mg grain pellet; Bio-Serv, Flemington, NJ, USA) and syringe pump through which 20% sucrose/10% polycose solution (0.2ml) could be delivered. An infrared beam crossed the magazine to detect head entries. Each chamber was equipped with two retractable levers that were located to the left or right of the magazine. The chambers were also fitted with a white-noise generator, a sonalert that delivered a 3 kHz tone, and a sonaloid that delivered a 5Hz clicker when activated. All stimuli were adjusted to 80dB in the presence of background noise (∼60dB) provided by the ventilation fan fixed to the cabinet throughout training and testing. A house light (3W, 24V) located on the opposite wall to the magazine provided constant illumination. Two microcomputers running the Med-PC program (Med Associates) were used to program and control all experimental events and record experimental data. Outcome devaluation prefeeding procedures were conducted in transparent plastic boxes with wire top lids.

#### Surgery

Stereotaxic surgery was conducted under isoflurane anaesthesia (5% induction; 1-2% maintenance). Rats were placed in a stereotaxic frame (Kopf Instruments, Tujunga, CA, USA), after which bupivacaine hydrochloride (0.1ml) was injected subcutaneously at the incision site. An incision was made into the scalp to expose the skull surface, and the incisor bar was adjusted to align bregma and lambda on the same horizontal plane. Rats received infusions into the NAC core at the following coordinates (in mm relative to bregma): anteroposterior +1.4, mediolateral ±2.2, dorsoventral -7.5. 1µl of pAAV-GFAP-hM4D(Gi)-mCherry (#50479; Addgene, Watertown, MA, USA) was infused at a rate of 0.2µl/min in each hemisphere. The needle was left in place for two min prior to removal to allow for diffusion. Control animals received an identical procedure except they were infused in the NAC core with pAAV-GFAP104-mCherry (#58909; Addgene) instead. All rats then received a subcutaneous injection of 0.1ml metacam (CenVet). Rats were given seven days to recover from surgery, after which they received three days of food restriction prior to the commencement of behavioural procedures. Rats were weighed and handled daily during this period.

#### Behavioural procedures

##### Pavlovian training

Rats first received eight days of Pavlovian training in which two conditioned stimuli (white noise or clicker) were presented for two min and each paired with the delivery of one of the two food outcomes (pellets or sucrose) on a random time 30 s schedule throughout the stimulus duration. Each stimulus was presented four times across the 60 min session in a pseudorandom order with a variable intertrial interval (ITI) averaging five min. Stimulus-outcome (S-O) pairings were counterbalanced so that half of the rats in each group received noise-pellet, clicker-sucrose contingencies, and the remaining half received the opposite arrangement. Magazine entries were recorded throughout the session and analysed by comparing responses during the two min stimulus duration to the two min prior to the stimulus presentation (Pre-S).

##### Instrumental training

After Pavlovian training, rats received eight days of lever press training in which a left or right lever press could be performed to obtain one of the two food outcomes. Each session lasted for a maximum of 50 min and consisted of two 10 min sessions on each lever (i.e. four x 10min sessions in total) separated by a 2.5 min time-out period in which the levers were retracted, and the house light turned off. If animals earned more than 20 outcomes on one lever during a 10min session, the session terminated, the lever retracted and the house light turned off, and the 2.5min time-out period initiated. Rats could earn a maximum of 80 outcomes per session (40 pellets, 40 sucrose). Action-outcome (A-O) pairings were counterbalanced so that half of the rats in each group received left lever-pellet, right lever-sucrose, and the remaining half received the opposite arrangement. Lever pressing was continuously reinforced on the first two days. Rats were then shifted to a random ratio (RR)-5 schedule for the next three days (i.e. each action delivered an outcome with a probability of 0.2), then to an RR10 schedule (i.e. outcome probability of 0.1) for the last three days.

##### Specific Pavlovian-instrumental transfer

Following the last day of instrumental training, rats were tested for specific Pavlovian-instrumental transfer. Approximately 25min prior to testing, rats received an i.p. injection of Deschloroclozapine (DCZ; National Institute of Mental Health) or vehicle (VEH; saline). DCZ was dissolved in 0.9% saline and administered at a dose of 0.1mg/kg for all behavioural testing. Throughout the test session, both levers were extended and no outcomes were delivered (i.e. conducted in extinction). Baseline responding was first extinguished for seven min prior to any stimuli being presented. Each stimulus was then presented four times for two min each across 40min in the following order: noise-clicker-clicker-noise-clicker-noise-noise-clicker. Each stimulus presentation was separated by a fixed three min ITI. Magazine entries and lever press rates were recorded throughout the session.

##### Outcome devaluation

The day following specific transfer, rats received one day of instrumental retraining on RR10 as previously described. Following this session, rats were put into the devaluation boxes for 1 hr with a small amount of their daily chow allowance to habituate them to the boxes.

One day later, rats received their first outcome devaluation test. They received an i.p. injection of DCZ or vehicle and then immediately received free access to one of the two outcomes (20g pellets or 200ml sucrose) in the devaluation box for 40-45min. Rats were then placed in the operant chambers for a 10min outcome devaluation test. During this test, both levers were extended and lever presses were recorded, but no outcomes delivered. The following day, the rats received a second outcome devaluation test in which the alternative food outcome was freely provided prior to testing. The order in which outcomes were devalued were counterbalanced within- and between-groups. Results are reported as an average across the two outcome devaluation tests.

##### Outcome-selective reinstatement

The day following outcome devaluation testing, rats underwent an outcome-selective reinstatement test. Twenty-five min prior to this test, rats received an i.p. injection of DCZ or vehicle. The test session began with a three min extinction period to lower baseline responding on both levers. Rats then received four reinstatement trials with an ITI of four min. Each reinstatement trial consisted of a single delivery of one of the two outcomes (pellet or sucrose) in the following order: sucrose-pellet-pellet-sucrose. Responding was measured during the two min periods immediately before (Pre) and after (Post) outcome delivery.

##### Tissue preparation and microscopy

In the 2-3 days following behavioural testing, animals were euthanised via CO2 inhalation and transcardially perfused with 4% paraformaldehyde (PFA) in 0.1M sodium phosphate buffer (PBS; pH 7.3-7.5). The brains were removed and postfixed in 4% PFA overnight then placed in 30% sucrose until they sank. Brains were sectioned coronally at 40µm through the NAC core using a cryostat (NX70; Leica Microsystems, North Ryde, NSW, Australia) maintained at approximately -20°C and the sections were stored in cryoprotectant solution at -20°C. To assess placement, three representative sections were selected for each rat. The sections were washed in PBS three times (10min per wash) and then incubated in a blocking solution comprising of 3% Bovine Serum Albumin (BSA), 0.25% TritonX-100 in PBS for 2h. Sections were then incubated for 48h in a blocking solution containing mouse anti-GFAP (1:300; #3670, Cell Signaling Technology, Danvers, MA, USA), rabbit anti-DSred (1:1000; #632496, Takara, Shiga, Japan), and chicken anti-NeuN (1:1000; #GTX00837, GeneTex, Irvine, CA, USA) diluted in PBS containing 3% BSA and 0.25% TritonX-100 at room temperature. The sections were then washed three times in PBS (10 min per wash). They were then incubated in a blocking solution overnight containing goat anti-mouse AF488 (1:500; #A-11001, Thermo Fisher Scientific, North Ryde, NSW, Australia), donkey anti-rabbit AF568 (1:500; #A10042, Thermo Fisher Scientific), and goat anti-chicken Dylight755 (1:500; #SA5-10075, Thermo Fisher Scientific) in PBS containing 3% BSA and 0.25% TritonX-100 at room temperature.

Sections were washed three times in PBS (10min per wash) and were then counterstained with DAPI (1:1000; Thermo Fisher Scientific) diluted in PBS for 10-15min. Sections received three final washes (10min per wash) prior to being mounted onto Superfrost microscope slides (Thermo Fisher Scientific) using Vectashield Antifade Mounting Medium (#H-1000; Vector Laboratories, Newark, CA, USA). Imaging was conducted on a slide scanner (Zeiss Axioscan Z1Slide Scanning Microscope, Macquarie Park, NSW, Australia). Subjects whose viral infection was minimal or misplaced were excluded from the statistical analysis.

##### Data analyses

All data were analysed using planned orthogonal contrasts controlling the per-contrast error rate at α = 0.05, using the method described by Hays (Hays, 1973). Training data were analysed using contrasts testing for linear trends, and interactions with group, day, and/or stimulus period (where relevant). Analyses of test data first established that there were no overall differences (i.e. main effects) in responding between the control groups, hM4Di+VEH and mCherry+DCZ, followed by comparing these groups (averaged) to group hM4Di+DCZ. Outcome devaluation test data was reported in both raw form and as a percentage of baseline responding, with baseline defined as the average left and right lever presses on the two days of instrumental training immediately preceding the test. Significant interactions were followed by simple-effects analyses.

Data are presented averaged across counterbalanced conditions.

### Experiment 2. The effect of NAC core astrocyte disruption on general PIT

#### Animals

A total of 37 Long Evans rats (18 female and 19 male) were used as subjects in this experiment and were as described in Experiment 1. Three groups were generated: hM4Di+VEH (*n* = 11, 5 female, 6 male), mCherry+DCZ (*n* = 14, 7 female, 7 male), and hM4Di+DCZ (*n* = 12, 6 female, 6 male).

#### Apparatus

All apparatus were as described in Experiment 1.

#### Surgery

All surgery procedures were as outlined in Experiment 1.

#### Behavioural procedures

##### Pavlovian training

Rats received eight days of Pavlovian training in which a single stimulus (S+, white noise or clicker) was presented for two min and paired with the delivery of a food outcome (pellets or sucrose) on a random time 30s schedule throughout the stimulus duration. The stimulus was presented six times across the 45min session with a variable ITI averaging five min. stimulus-outcome pairings were counterbalanced such that an equal number of rats received one of the following pairings: noise-pellets, noise-sucrose, clicker-pellets, or clicker-sucrose. Magazine entries were recorded throughout the session and analysed by comparing responses during the two min stimulus compared to the two min prior to the stimulus presentation (Pre-S).

##### Instrumental training

Rats next received eight days of instrumental training in which a lever press on a single lever (left or right) could be made to obtain the alternate food outcome (e.g. if rats received pellets during Pavlovian training, they received sucrose during instrumental training). The session was terminated after 30min had elapsed or animals had earned 30 outcomes. Action-outcome pairings were counterbalanced such that equal numbers of rats in each group received one of the following contingencies: left lever-pellets, right lever-pellets, left lever-sucrose, or right lever-sucrose. Reinforcement schedules were as outlined in Experiment 1.

##### Pavlovian exposure to S-

After instrumental training, rats received one session where they were exposed to the alternate stimulus (S-, i.e. if they had received Pavlovian training with the noise this was the clicker, and vice versa) with a variable ITI averaging five min with no outcomes delivered.

##### General Pavlovian-instrumental transfer

The day following Pavlovian exposure to the S-, rats were tested for general Pavlovian-instrumental transfer. Rats received an i.p. injection of DCZ or vehicle (saline) 25 min prior to test. During the test session, the lever trained during instrumental training was continuously available, however lever presses did not earn any food outcomes. Baseline responding was extinguished for seven min prior to any stimulus being presented. Across the 40min test, both the S+ and S– were presented four times, each presentation lasting two min in the following order: noise-clicker-clicker-noise-clicker-noise-noise-clicker. Each stimulus presentation was separated by a fixed three min ITI. Magazine entries and lever press rates were recorded throughout the session.

##### Tissue preparation and microscopy

General tissue and microscopy procedures were as described in Experiment 1. For assessment of placements, sections were incubated overnight at room temperature in a blocking solution containing rabbit anti-DSRED (1:1000; #632496, Takara, Shiga, Japan). They were then incubated for 2h at room temperature in a blocking solution containing donkey anti-rabbit AF568 (1:500; #A10042, Thermo Fisher Scientific). Subjects whose viral infection was minimal or misplaced were excluded from the statistical analysis of behavioural tasks. For quantification of phospho-pyruvate dehydrogenase (pPDH), a subset of animals (hM4Di+VEH [*n* = 4], mCherry+DCZ [*n* = 4], and hM4Di+DCZ [*n* = 5]) were transcardially perfused 45min post-DCZ injection. pPDH is a marker for neuronal inhibition, as it inversely correlates with the intensity of action potential firing in cells (Yang et al., 2024). Four representative sections containing the target NAC core were collected for each animal and incubated for 72h at 4°C in a blocking solution containing chicken anti-NeuN (1:500; #GTX00837, GeneTex) and rabbit anti-pPDH (1:1000; #37115S, Cell Signaling Technology). Sections were then incubated for 2h in a blocking solution containing goat anti-rabbit 488 (1:500; #A11008, Thermo Fisher Scientific) and goat anti-chicken 647 (1:400; #A-21449, Thermo Fisher Scientific). Z-stack images for quantification of co-localisation was conducted on a confocal microscope (20X, STELLARIS 8, Leica Microsystems, Lane Cove West, Australia) to enable quantification of a high number of cells per region while balancing for high somatic resolution. The auto-mCherry channel, indicating viral transfection, was used to determine the best confocal plane to acquire the image.

##### Image quantification

Immunofluorescence analysis was conducted using the Fiji ImageJ software (Schindelin et al., 2012). The pPDH and NeuN channels were separated for individual analysis and z-sections were summed together. A mask was drawn around individual pPDH-positive cells or neurons using thresholds. Mean gray value, ROI count and size, and ROI morphology (perimeter, circularity) were measured in each channel. These values were averaged across the four bilateral slices per animal to calculate mean values per measure, and these means were used in statistical analyses.

##### Data analyses

All behavioural data was analysed as described in Experiment 1. Quantification analyses first established that there were no significant differences between the control groups, hM4Di+VEH and mCherry+DCZ, which was then followed by comparing these control groups to the group hM4Di+DCZ.

## Results

### Experiment 1: Physiological astrocytic signalling in NAC core is necessary for goal-directed responding but not for cue- or outcome-guided choice

The aim of Experiment 1 was to investigate whether astrocytes in the nucleus accumbens (NAC) core contribute to various forms of choice behaviour for food outcomes. Rats first received bilateral injections of an adenovirus (AAV) that caused astrocytes to express the inhibitory hM4Di designer receptor exclusively activated by a designer drug (DREADD), or a control virus that led to astrocyte-specific expression of the fluorophore. Placements and transfections are shown in Figure 1A. Following removal of animals with misplaced or lack of transfection final numbers were: hM4Di+VEH (*n* = 10, 5 female, 5 male), mCherry+DCZ (*n* = 10, 5 female, 5 male), and hM4Di+DCZ (*n* = 10, 4 female, 6 male).

**Figure 1.**
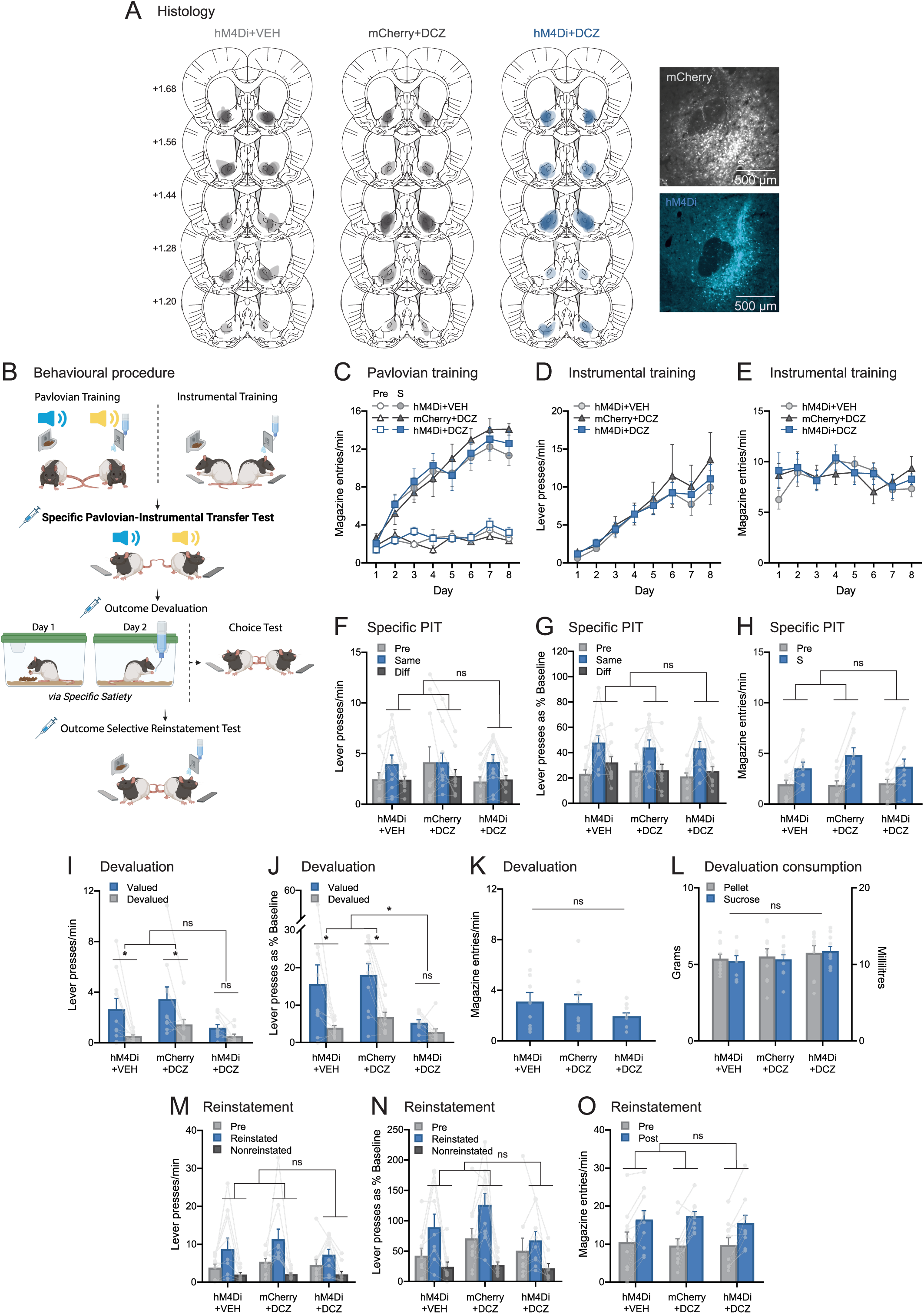
Chemogenetic activation of the Gi-pathway in NAC core astrocytes disrupts outcome devaluation but not outcome-specific PIT or reinstatement. (**A**) Diagrammatical representation of the location and extent of NAC core viral infections included in the analyses for hM4Di+VEH (n = 10, 5 female, 5 male), mCherry+DCZ (n = 10, 5 female, 5 male), and hM4Di+DCZ (n = 10, 4 female, 6 male). Distances are indicated in mm from bregma. Micrographs showing viral expression in the NAC core for mCherry (top right) and hM4Di (bottom right). (**B**) Behavioural procedures for outcome-specific Pavlovian-instrumental transfer, outcome devaluation, and outcome-selective reinstatement, created with Biorender. (**C**) Magazine entries per min during Pavlovian training. Lever presses per min (**D**) and magazine entries per min (**E**) during instrumental training. (**F**-**H**) Outcome-specific PIT: lever presses per min (**F**), lever presses as a percentage of baseline responding (**G**), and magazine entries per min (**H**). (**I**-**L**) Outcome devaluation: lever presses per min (**I**), lever presses as a percentage of baseline responding (**J**), magazine entries per min (**K**), and outcome consumption prior to test (**L**). (**M**-**O**) Outcome selective reinstatement: lever presses per min (**M**), lever presses as a percentage of baseline responding (**N**), and magazine entries per min (**O**). Data are shown as mean ± SEM. Panels **F**-**O** include individual data points for each rat. Asterisk (*) denotes p < 0.05, ns denotes non-significant.

The design of this experiment is shown in Figure 1B. Following recovery from surgery and a short period of food deprivation, rats received Pavlovian training followed by instrumental training, both of which were conducted drug-free. Over the following days, rats were given injections of DCZ (groups mCherry+DCZ and hM4Di+DCZ) or vehicle (group hM4Di+VEH) and tested on specific Pavlovian-instrumental transfer, outcome devaluation, and outcome-selective reinstatement. Given prior findings that lesions (Corbit and Balleine, 2011; Corbit et al., 2016) and neuroinflammation (Abiero et al., 2025) in the NAC core leave specific transfer and selective reinstatement intact, but impair or attenuate outcome devaluation, we expected that disrupting astrocytic signalling in NAC core astrocytes would likewise affect devaluation but not transfer or reinstatement. We further predicted that magazine entries would be affected by this manipulation, in light of evidence that NAC core lesions reduce this response (Balleine and Killcross, 1994) and NAC core neuroinflammation increases it (Abiero et al., 2025).

As expected, Pavlovian and instrumental acquisition did not differ between groups. Magazine entries during Pavlovian conditioning are shown in Figure 1C. Rats in all groups learned the S-O relationships, as they made progressively more magazine entries per minute during stimulus presentations than the 2 min pre-S over days. This is supported by a linear main effect, F(1, 27) = 128.73, p < 0.001, and no main effects of group and no group x linear interactions, all Fs < 1. There was, however, a main effect of stimulus period (S > pre-S), F(1,27) = 229.91, p < 0.001, and a linear x stimulus period interaction, F(1,24) = 171.95 p < 0.001, neither of which interacted with group comparisons, all Fs < 1.

Likewise, during instrumental training, rats in each group successfully learned to press left and right levers for the sucrose and pellet outcomes (Fig. 1D). There was again a linear main effect, F(1,23) = 47.71, p < .001 (note that data for 4 rats is missing from this analysis due to a MedPC error – 1 each from groups hM4Di+VEH and hM4Di+DCZ and 2 from mCherry+DCZ) that did not differ according to group; largest F was for the hM4Di+VEH vs. mCherry+DCZ x linear interaction, F(1,23) = 1.28, p = 0.27. Magazine entries did not differ significantly between groups during instrumental conditioning, all Fs < 1 (Fig. 1E).

Rats were next tested on specific Pavlovian-instrumental transfer. As expected, rats in all groups pressed the lever associated with the same outcome as the currently presented stimulus more than the different lever (i.e. the pellet stimulus elicited pressing on the pellet lever, and the sucrose stimulus on the sucrose lever: Same > Different, Fig. 1F). There were no group main effects, both Fs < 1, but there was a main effect of stimulus (Same > Different), F(1,27) = 19.1, p < 0.001, that did not interact with either group comparison, Fs < 1. Calculating test data as a percentage of baseline did not alter results, all group main effects and interactions, Fs < 1 (Fig. 1G). As shown in Figure 1H, magazine entries also did not differ between groups, main effects of group, Fs < 1, main effect of stimulus period (S > pre-S), F(1,27) = 33.87, p < 0.001. The largest F involving a group difference was for the hM4Di+VEH vs. mCherry+DCZ x stimulus period interaction, F(1,27) = 2.6, p = 0.118.

Following specific transfer, rats underwent outcome devaluation tests to assess goal-directed control of behaviour, as shown in Figure 1I. Devaluation was expected to be intact for controls (Valued > Devalued) and impaired for group hM4Di+DCZ (Valued = Devalued). Results were largely as expected, because overall responding (i.e. responding averaged across valued and devalued levers) was marginally reduced in group hM4Di+DCZ relative to controls, F(1,27) = 3.85, p = 0.06, whereas responding did not differ between the two control groups, F(1,27) = 1.54, p = 0.225.

The selectivity of responding on the Valued relative to the Devalued lever was also attenuated in group hM4Di+DCZ relative to the controls. There was a main effect of devaluation, F(1,27) = 19.64, p < 0.001, that did not interact with the comparison between control groups, F < 1. The control group vs. hM4Di+DCZ x devaluation interaction also did not reach significance, F(1,27) = 3.317, p = .079, however there were significant simple effects (Valued > Devalued) for each control group; F(1,27) = 11.55, p = 0.002, for hM4Di+VEH and F(1,27) = 10.28, p = 0.003, for group mCherry+DCZ, and no such effect for group hM4Di+DCZ (Valued = Devalued), F(1,27) = 1.15, p = 0.29.

The failure of group main effects and interactions to reach significance appeared to be the result of subtle baseline differences, as group differences were accentuated when test data was calculated as a percentage of baseline responding (Fig. 1J). Specifically, the main effect of group (hM4Di+DCZ vs. controls) was now significant, hM4Di+DCZ, F(1,27) = 8.47, p =0 .007, and this interacted with devaluation, F(1,27) = 4.82, p = 0.037. Importantly, the comparison between the two control groups was still non-significant, main effect and interaction Fs < 1. This pattern was again supported by significant simple effects for both control groups, hM4Di+VEH, F(1,27) = 12.03, p = 0.002, and mCherry+DCZ, F(1,27) = 11.38, p = 0.002, but not group hM4Di+DCZ, F < 1. Surprisingly, magazine entries did not differ significantly between groups during the devaluation test, either for the comparison between the two control groups, F < 1, or for the hM4Di+DCZ vs. control groups comparison, F(1,27) = 2.16, p = 0.153 (Fig. 1K).

Finally, as predicted during outcome-selective reinstatement tests, performance was intact (Reinstated > NonReinstated) for all groups (Fig. 1M). There were no main effects of group, Fs < 1, but there was a main effect of reinstatement, F(1,27) = 31.08, p < 0.001, that did not interact with either the control group comparison, F < 1, or the control groups versus group hM4Di+DCZ comparison, F(1,27) = 1.11, p = 0.301. Calculating data as a percentage of baseline did not alter these findings, since the largest F for these adjusted data was for the group x reinstatement interaction, F(1,26) = 2.58, p = 0.12 (Fig. 1N). Once again, magazine entries did not differ significantly between groups on this test, with Fs < 1 for all main effects and interactions (Fig. 1O).

Together, these data show that the disruption of astrocytic signalling in the NAC core impairs goal-directed responding in a free operant situation but leaves selective choice intact when it is guided by cues that signal specific outcomes, or by the outcomes themselves. They further show that, against expectations, head entry responses into the food magazine do not appear to rely on astrocytic signalling in NAC core in any of the tested paradigms.

### Experiment 2: General Pavlovian instrumental transfer relies on intact astrocytic signalling in the NAC core

The results of Experiment 1 indicate that intact NAC core astrocyte signalling is necessary for the influence of general arousal or motivation over instrumental actions, but not for the selection of those same actions when they are guided by cues – regardless of whether those cues are outcome associated stimuli (in specific transfer) or the outcomes themselves (in reinstatement). It is therefore curious as to why studies have found that drug-associated cues do appear to influence instrumental actions in a manner that is dependent on NAC core astrocyte function. That is, several studies have found that, in animals taught to self-administer cocaine (Scofield et al., 2015, 2016) or opioids (Kruyer et al., 2019, 2022), the disruption of NAC core astrocyte function prevents the reinstatement of this responding when animals are exposed to the cues that signal them. However, these findings seem to be specific to cues signalling drug outcomes, as the same studies do not find similar results when using food rewards such as sucrose.

This raises the question of whether cues signalling drug but not food outcomes engage NAC core astrocytes due to differences in their inherent sensory-specific properties, or in the pharmacological changes they create, or because they are just generally more arousing. A study by Corbit et al., (2016) suggests that it could be the latter, because they showed that whereas sucrose-cues only elicited responding on a lever associated with sucrose, alcohol-cues elicited responding on levers associated with either alcohol or sucrose, and only the effect of (the generally arousing) alcohol cues on responding depended on the integrity of the NAC core. Although this study did not investigate the role of astrocytes specifically, if we extrapolate from these studies that NAC core astrocyte signalling might be similarly necessary for the ability of generally arousing cues to elevate responding, then disrupting their function would be also expected to impair general Pavlovian instrumental responding using food outcomes. This was tested in Experiment 2, the design for which is shown in Figure 2B. Following the removal of rats with misplaced or no signs of transfection, final group numbers for this experiment were: hM4Di+VEH (*n* = 6, 1 female, 5 male), mCherry+DCZ (*n* = 13, 7 female, 6 male), and hM4Di+DCZ (*n* = 8, 4 female, 4 male).

**Figure 2.**
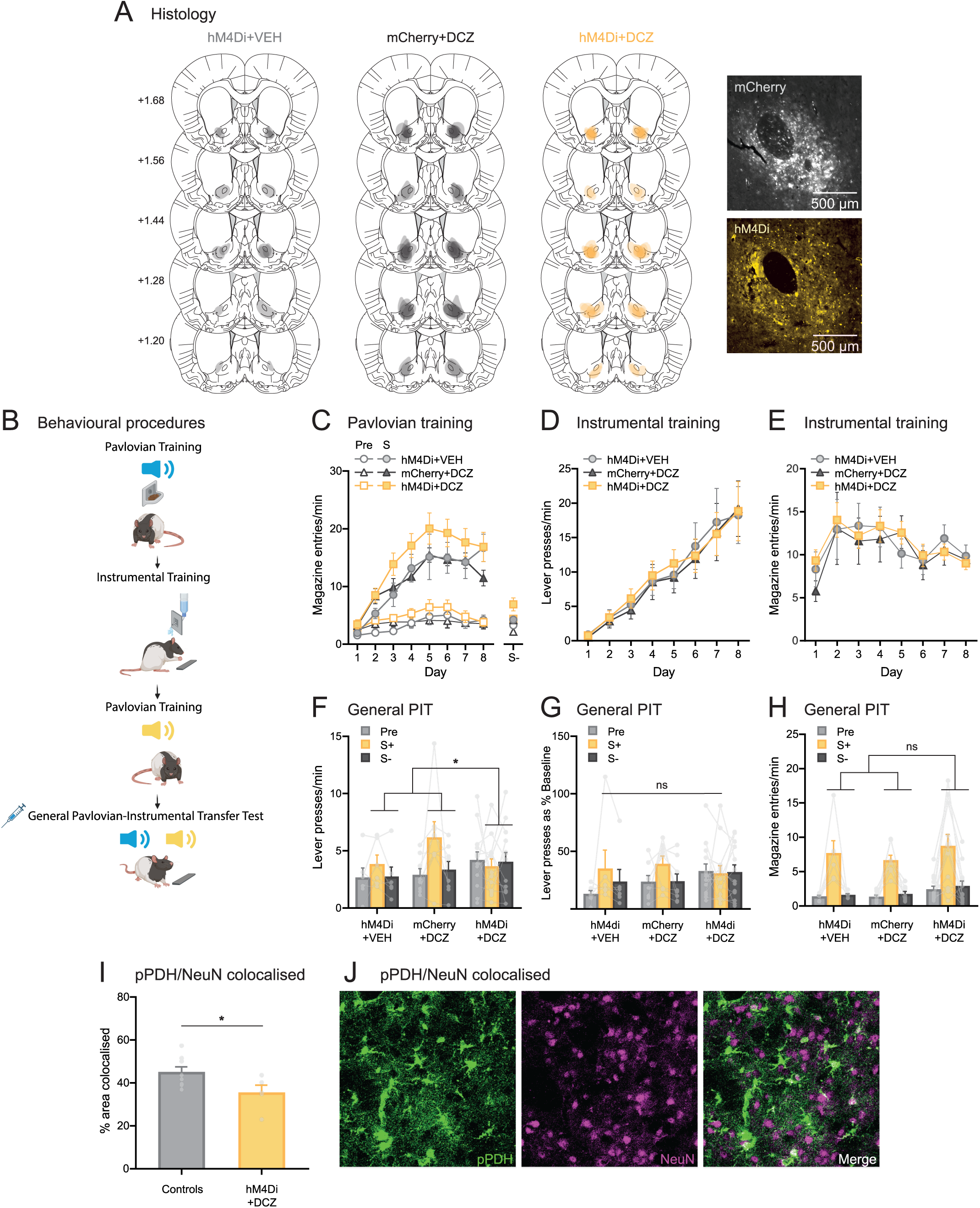
Chemogenetic activation of the Gi-pathway in NAC core astrocytes disrupts general PIT. **(A)** Diagrammatical representation of the location and extent of NAC core viral infections included in the analyses for hM4Di+VEH (n = 6, 1 female, 5 male), mCherry+DCZ (n = 13, 7 female, 6 male), and hM4Di+DCZ (n = 8, 4 female, 4 male). Distances are indicated in mm from bregma. Micrographs showing viral expression in the NAC core for mCherry (top right) and hM4Di (bottom right). (**B**) Behavioural procedures for general Pavlovian-instrumental transfer, created with Biorender. (**C**) Magazine entries per min during Pavlovian training. Lever presses per min (**D**) and magazine entries per min (**E**) during instrumental training. (**F**-**H**) General PIT: lever presses per min (**F**), lever presses as a percentage of baseline responding (**G**), and magazine entries per min (**H**). (**I**) Percentage of area showing colocalisation between pPDH and NeuN averaged across controls (hM4Di+VEH [n = 4] and mCherry+DCZ [n = 4]) and hM4Di+DCZ (n = 5). (**J**) Representative images showing colocalisation of pPDH with NeuN. Data are shown as mean ± SEM. Panels **F**-**I** include individual data points for each rat. Asterisk (*) denotes p < 0.05, ns denotes non-significant.

Rats first received Pavlovian training which consisted of presentations of a single S+ associated with pellets or sucrose, followed by instrumental training in which they learned to press a single lever (left or right) for the alternate outcome. Rats then received a single day of training with an S-stimulus paired with nothing. Behavioural training was conducted drug-free, where rats received injections of DCZ or vehicle 25 min prior to the general PIT test. If undisrupted homeostatic astrocytic signalling was required for the generally arousing effects of cues on actions, then the control groups should show intact general PIT (i.e. S+ > S-/baseline) whereas this should be impaired for group hM4Di+DCZ (S+ = S-/baseline).

Following the completion of behavioural training and testing, we wished to determine the cellular effect of activating the Gi pathway in NAC core by performing perfusions 45 min after DCZ or vehicle injection (i.e. 20 min for DCZ to take effect, and 25 min for peak expression of the immunomarker), and measuring levels of pPDH in neurons, a marker that has been called the ‘inverse of c-Fos’ in that its presence indicates neuronal inhibition rather than activation (Yang et al., 2024).

Responding during Pavlovian training is shown in Figure 2C. As shown, rats in each group acquired conditioning to the S+ because they made progressively more magazine entries during the stimulus relative to the 2 min pre-S period over days. There were no group main effects (F < 1 for the comparison between control groups), although the Control groups versus hM4Di+DCZ comparison could be considered marginal, F(1,24) = 3.66, p = 0.068. Importantly, however, the direction of this difference is that of *higher* responding in the hM4Di+DCZ group relative to controls which, if anything, might be expected to facilitate rather than impair general PIT. Moreover, the rate of acquisition did not differ between groups because there was a linear x stimulus period (pre-S vs. S) interaction, F(1,24) = 71.09, p < 0.001, that did not interact with any group comparisons, all Fs < 1. As shown in Figure 2D, there were also no group differences during instrumental lever press acquisition, as there were no main effects of group, Fs < 1, and a linear main effect, F(1,24) = 51.6, p < 0.001, that did not interact with any group comparisons, Fs < 1. Magazine entries were also similar between groups during instrumental conditioning, all Fs < 1 (Fig. 2E).

Data from the general PIT test are shown in Figure 2F. General PIT was intact for rats in both the hM4Di+VEH and mCherry+DCZ control groups, who pressed the lever at higher rates during S+ presentations compared to responding at baseline or during presentation of S-. In contrast, general PIT was impaired for group hM4Di+DCZ whose responding remained flat across all three time periods. There were no main effects of group, largest F for the control group comparison, F(1,24) = 1.01, p = 0.325, but there was a marginal main effect of higher responding during S+ presentations than at other time periods, F(1,24) = 3.69, p = 0.067. Importantly, whereas this difference (i.e. S+ > baseline/S-) did not interact with the comparison between control groups, F(1,24) = 1.28, p = 0.269, it did significantly interact with the control group versus hM4Di+DCZ comparison, F(1,24) = 4.35, p = 0.048. To be consistent with Experiment 1 results, we have also presented the data as a percentage of baseline responding, however, although this preserved the directional effects, in this case it did cause the interaction to no longer be significant (F(1,24) = 3.05, p = .094, Fig. 2G).

This impairment in general PIT was specific to the instrumental lever press response, because as shown in Figure 2H, rats in all groups made more magazine entries during S+ presentations relative to other time periods. There was a main effect showing elevated responding during S+ relative to baseline and S-presentations, F(1,24) = 37.38, p < 0.001, no group main effects (largest F for the control versus hM4Di+DCZ comparison, F(1,24) = 2.25, p = 0.147), and no group x stimulus period interactions, Fs < 1. Together, these data support the prediction that intact astrocytic signalling in NAC core is necessary for the generally arousing effect reward-predictive cues have on instrumental responding specifically.

Finally, NAC core levels of pPDH for a subset of animals from each group are shown in Figure 2I. Because only 4 animals from each control group contributed to this subset, we collapsed across them (n = 8) to compare to group hM4Di+DCZ (n = 5). The percentage of area of pPDH co-localised with NeuN was significantly higher in controls than in group hM4Di+DCZ, suggesting that the controls experienced more neuronal inhibition. This result suggests that chemogenetic activation of the Gi-pathway in NAC core astrocytes increased the activity of nearby neurons, which is likely to have contributed to the behavioural disruptions observed.

## Discussion

Current results demonstrate a role for NAC core astrocytes in modulating instrumental responding when it is driven by general arousal or motivation but not by specific outcome properties. In Experiment 1, chemogenetic activation of Gi-coupled receptors on NAC core astrocytes impaired the performance of outcome devaluation relative to controls, whereas cued choice behaviours were intact despite this manipulation, both in specific PIT and in outcome-selective reinstatement. In Experiment 2, the same manipulation prevented cue-elicited instrumental responding in a general PIT paradigm, indicating that undisrupted astrocyte function in the NAC core is necessary for instrumental responding to cues when those cues are generally arousing. These findings extend previous evidence that NAC core astrocytes regulate instrumental responding, revealing that their role is not limited to drug-seeking behaviours but also applies to actions invigorated by arousal states more generally.

Disrupting the signalling of NAC core astrocytes here produced alterations in instrumental responding that are largely consistent with the consequences of NAC core lesions seen in previous studies. In particular, prior studies have demonstrated that excitotoxic lesions of this brain region also impair outcome devaluation (Corbit et al., 2001) and general transfer (Corbit and Balleine, 2011; Corbit et al., 2016) whilst leaving specific PIT intact. However, one notable difference between current findings and the effects of NAC core lesions is that we found no effect of NAC core astrocyte manipulations on magazine entries in any phase of either experiment, whereas Balleine and Killcross (1994) reported a reduction in magazine entries following ibotenic lesions of NAC core, particularly when rats had undergone a shift in incentive value or deprivation state. Thus, just as we observed in the posterior dorsomedial striatum (Abiero et al., 2025), activating the Gi-pathway in astrocytes in the NAC core produces behavioural effects that are somewhat dissociable from the inactivation of neurons, highlighting the unique contribution of astrocytes to decision-making behaviours.

The behavioural effects reported here further support the notion that astrocytic function is highly region-specific. This is because, in contrast to the current finding that specific PIT does not rely on NAC core astrocytes, disrupting the signalling of astrocytes in the pDMS was shown to impair specific PIT (Abiero et al., 2025). This dissociation is particularly striking given the relatively homogeneous cellular architecture of the dorsal and ventral striatum, both of which consist of more than 95% medium spiny neurons, with approximately half expressing D1-receptors and half expressing D2 receptors (Beckstead, 1988; Gerfen et al., 1990; Matamales et al., 2020). The remaining population comprising of cholinergic, fast-spiking, and other types of interneurons (Luk and Sadikot, 2001; Ding et al., 2010). However, whether the regional differences in the roles of astrocytes in each region stem from intrinsic differences in the astrocyte physiology, or from region-specific astrocyte-neuron interactions (Ganesan et al., 2025), remains to be determined. Notably, the activation of Gi-coupled receptors on astrocytes appears to have increased neuronal activation in both regions, as shown in the pDMS using electrophysiology (Abiero et al., 2025) and here in the NAC core using immunohistochemistry. However, the use of different measurement techniques prevents direct comparison and future studies are required to identify the precise cellular mechanisms that give rise to these distinct behavioural outcomes.

Nevertheless, the existing literature does provide some insight into the NAC core mechanisms that are unlikel*y* to underlie actions elicited by generally motivating cues. Work by Kalivas and colleagues (Scofield et al., 2016; Bobadilla et al., 2017; Kruyer et al., 2019, 2022) has uncovered several lines of evidence indicating that an increase in sensitivity to drug cues is underpinned by increases in extracellular glutamate in NAC core that occur as a result of reduced astroglial glutamatergic transport. However, this mechanism cannot account for present findings, as the pPDH data revealed higher levels of neuronal *inhibition* in the control groups despite these animals displaying greater cue sensitivity than group hM4Di+DCZ. Thus, although responses to both drug cues and cues that have generally motivating properties depend on NAC core astrocytes, the mechanism by which they do so is non-identical.

Likewise, the mechanism that proposed to underlie NAC core-dependent cue-induced sucrose seeking also fails to explain current results. Specifically, Bobadilla et al., (2017) suggested that this depends on alterations to presynaptic metabotropic glutamate receptor2/3 (mGluR2/3) signalling. However, because this mechanism is astrocyte-independent, it cannot account for why disrupting astrocyte signalling in the current study affected responding driven by alterations in incentive value. Taken together, therefore, current and prior findings suggest that the cellular mechanisms within NAC core that underpin cue-driven behaviour vary depending on what is being signalled by the cue (e.g. food or drugs), and the motivational process it engages (seeking specific outcomes versus general arousal).

Of course, there are potential mechanisms outside of glutamate transport that are also affected by astrocyte manipulations that could have produced the current behavioural effects. For instance, one recent study (Corkrum et al., 2020) found that G-protein coupled receptor signalling in astrocytes was necessary for the dopaminergic depression of excitatory post-synaptic currents (EPSCs) in the NAC core. Although that study used Gq-DREADDs (as opposed to the Gi-DREADDs used in the current study) current manipulations would likely also have altered the physiological occurrences of dopamine-depressing EPSCs, which could have led to the observed reductions in responding under generally arousing states. Again, this will need to be determined by future studies.

A final complexity to consider is that, in contrast to current findings, we have previously shown that neuroinflammation in the NAC core, induced by local injections of lipopolysaccharide (LPS), increased the rate of magazine entries (Abiero et al., 2025), whereas no effect on magazine entries was observed in the current study. This is likely due to the differences in methodologies between the two studies, particularly the use of LPS-induced neuroinflammation versus astrocyte-targeted chemogenetics here. Indeed, although neuroinflammation in our prior study was found to alter the proliferation and phenotypic responses of astrocytes, it also affected microglia and likely triggered the release of various cytokines, among other possibilities. These broader inflammatory effects may account for the divergent behavioural outcomes. Moreover, neuroinflammation was present for the entirety of behavioural training, whereas the chemogenetic astrocytic disruptions occurred on test only.

Despite these remaining questions, the current data clearly show that the intact signalling properties of NAC core astrocytes are necessary for the performance of instrumental actions that are driven by the generally arousing properties of incentive value. Given the purported role of NAC core astrocytes in the development and maintenance of substance use disorder (Kitamura et al., 2010; Adermark and Bowers, 2016; Kruyer and Kalivas, 2021; Kruyer et al., 2022) and other compulsive phenotypes (Aida et al., 2015; Tanaka, 2021), these data suggests that their contribution may not lie in directing actions towards specific outcomes, but in enhancing the motivational drive behind those actions. It is important to note that this is just one of many physiological changes that occur in the brains of individuals with compulsive disorders and does not negate the possibility – or even probability – of parallel cognitive processes also contributing to symptoms. Indeed, taken together with prior research, the overall evidence is suggestive of several processes contributing to compulsion, between different individuals and within the same individual at different times.

## Acknowledgements

We thank the technical staff at the Ernst Facility at the University of Technology Sydney for support. The authors acknowledge the use of the Nikon TI (Nikon corporation),Stellaris 8 (Leica Microsystems) and Axioscan Z1 slide scanning (Zeiss) microscopes in the Microbial Imaging Facility in the Faculty of Science at the University of Technology Sydney. We would like to thank Dr Amy Bottomley & A/Prof Louise Cole for their technical assistance. Figure 2 was created with BioRender.com.

## Author contributions

L.A.B. and A.R.A. conceptualised and designed the research, J.M.G. and O.S., performed the research (i.e. data curation), L.A.B., J.M.G., M.D.K., and O.S., analysed the data, L.A.B., and K.M.T., acquired the funding, L.A.B., O.S., J.M.G., M.D.K., K.M.T. and A.R.A. wrote the paper (original draft – L.A.B. and O.S., review and editing – J.M.G., M.D.K., K.M.T., and A.R.A.)

## References

1. Abiero AR, Gladding JM, Iredale JA, Drury HR, Manning EE, Dayas CV, Dhungana A, Ganesan K, Turner K, Becchi S, Kendig MD, Nolan C, Balleine B, Castorina A, Cole L, Clemens KJ, Bradfield LA (2025) Dorsomedial striatal neuroinflammation causes excessive goal-directed action control by disrupting astrocyte function. Neuropsychopharmacol Available at: https://www.nature.com/articles/s41386-025-02247-4 [Accessed September 29, 2025].

2. Adermark L, Bowers MS (2016) Disentangling the Role of Astrocytes in Alcohol Use Disorder. Alcoholism Clin &amp; Exp Res 40:1802–1816.

3. Aida T, Yoshida J, Nomura M, Tanimura A, Iino Y, Soma M, Bai N, Ito Y, Cui W, Aizawa H, Yanagisawa M, Nagai T, Takata N, Tanaka KF, Takayanagi R, Kano M, Götz M, Hirase H, Tanaka K (2015) Astroglial Glutamate Transporter Deficiency Increases Synaptic Excitability and Leads to Pathological Repetitive Behaviors in Mice. Neuropsychopharmacol 40:1569–1579.

4. Balleine B, Killcross S (1994) Effects of ibotenic acid lesions of the nucleus accumbens on instrumental action. Behav Brain Res 65:181–193.

5. Beckstead RM (1988) Association of dopamine d, and d2 receptors with specific cellular elements in the basal ganglia of the cat: The uneven topography of dopamine receptors in the striatum is determined by intrinsic striatal cells, not nigrostriatal axons. Neuroscience 27:851–863.

6. Bobadilla A-C, Garcia-Keller C, Heinsbroek JA, Scofield MD, Chareunsouk V, Monforton C, Kalivas PW (2017) Accumbens Mechanisms for Cued Sucrose Seeking. Neuropsychopharmacology 42:2377–2386.

7. Chai H, Diaz-Castro B, Shigetomi E, Monte E, Octeau JC, Yu X, Cohn W, Rajendran PS, Vondriska TM, Whitelegge JP, Coppola G, Khakh BS (2017) Neural Circuit-Specialized Astrocytes: Transcriptomic, Proteomic, Morphological, and Functional Evidence. Neuron 95:531–549.e9.

8. Corbit LH, Balleine BW (2011) The general and outcome-specific forms of Pavlovian-instrumental transfer are differentially mediated by the nucleus accumbens core and shell. J Neurosci 31:11786–11794.

9. Corbit LH, Fischbach SC, Janak PH (2016) Nucleus accumbens core and shell are differentially involved in general and outcome-specific forms of Pavlovian-instrumental transfer with alcohol and sucrose rewards. Eur J Neurosci 43:1229–1236.

10. Corbit LH, Muir JL, Balleine BW (2001) The Role of the Nucleus Accumbens in Instrumental Conditioning: Evidence of a Functional Dissociation between Accumbens Core and Shell. J Neurosci 21:3251–3260.

11. Corkrum M, Covelo A, Lines J, Bellocchio L, Pisansky M, Loke K, Quintana R, Rothwell PE, Lujan R, Marsicano G, Martin ED, Thomas MJ, Kofuji P, Araque A (2020) Dopamine-Evoked Synaptic Regulation in the Nucleus Accumbens Requires Astrocyte Activity. Neuron 105:1036–1047.e5.

12. Ding JB, Guzman JN, Peterson JD, Goldberg JA, Surmeier DJ (2010) Thalamic Gating of Corticostriatal Signaling by Cholinergic Interneurons. Neuron 67:294–307.

13. Doya K (2008) Modulators of decision making. Nat Neurosci 11:410–416.

14. Durkee CA, Covelo A, Lines J, Kofuji P, Aguilar J, Araque A (2019) G i/o protein-coupled receptors inhibit neurons but activate astrocytes and stimulate gliotransmission. Glia 67:1076–1093.

15. Gerfen CR, Engber TM, Mahan LC, Susel Z, Chase TN, Monsma FJ, Sibley DR (1990) D1 and D2 Dopamine Receptor-regulated Gene Expression of Striatonigral and Striatopallidal Neurons. Science 250:1429–1432.

16. Hays, W. L. (1973) Statistics for the social sciences. New York, Holt, Rinehart, & Winston.

17. Kitamura O, Takeichi T, Wang EL, Tokunaga I, Ishigami A, Kubo S (2010) Microglial and astrocytic changes in the striatum of methamphetamine abusers. Legal Medicine 12:57–62.

18. Kruyer A, Angelis A, Garcia-Keller C, Li H, Kalivas PW (2022) Plasticity in astrocyte subpopulations regulates heroin relapse. Sci Adv 8:eabo7044.

19. Kruyer A, Kalivas PW (2021) Astrocytes as cellular mediators of cue reactivity in addiction. Curr Opin Pharmacol 56:1–6.

20. Kruyer A, Scofield MD, Wood D, Reissner KJ, Kalivas PW (2019) Heroin Cue-Evoked Astrocytic Structural Plasticity at Nucleus Accumbens Synapses Inhibits Heroin Seeking. Biol Psychiatry 86:811–819.

21. Luk KC, Sadikot AF (2001) GABA promotes survival but not proliferation of parvalbumin-immunoreactive interneurons in rodent neostriatum: an in vivo study with stereology. Neuroscience 104:93–103.

22. Matamales M, McGovern AE, Mi JD, Mazzone SB, Balleine BW, Bertran-Gonzalez J (2020) Local D2- to D1-neuron transmodulation updates goal-directed learning in the striatum. Science 367:549–555.

23. Schindelin J, Arganda-Carreras I, Frise E, Kaynig V, Longair M, Pietzsch T, Preibisch S, Rueden C, Saalfeld S, Schmid B, Tinevez J-Y, White DJ, Hartenstein V, Eliceiri K, Tomancak P, Cardona A (2012) Fiji: an open-source platform for biological-image analysis. Nat Methods 9:676–682.

24. Scofield MD, Boger HA, Smith RJ, Li H, Haydon PG, Kalivas PW (2015) Gq-DREADD Selectively Initiates Glial Glutamate Release and Inhibits Cue-induced Cocaine Seeking. Biol Psychiatry 78:441– 451.

25. Scofield MD, Li H, Siemsen BM, Healey KL, Tran PK, Woronoff N, Boger HA, Kalivas PW, Reissner KJ (2016) Cocaine Self-Administration and Extinction Leads to Reduced Glial Fibrillary Acidic Protein Expression and Morphometric Features of Astrocytes in the Nucleus Accumbens Core. Biol Psychiatry 80:207–215.

26. Tanaka K (2021) Astroglia and Obsessive Compulsive Disorder. In: Astrocytes in Psychiatric Disorders (Li B, Parpura V, Verkhratsky A, Scuderi C, eds), pp 139–149 Advances in Neurobiology. Cham: Springer International Publishing. Available at: https://link.springer.com/10.1007/978-3-030-77375-5_7 [Accessed June 24, 2025].

27. Yang D et al. (2024) Phosphorylation of pyruvate dehydrogenase inversely associates with neuronal activity. Neuron 112:959–971.e8.

